# Digit-Tracking Reveals Curiosity-Driven Visual Attention in Macaque Monkeys

**DOI:** 10.1101/2025.09.17.676830

**Authors:** Yidong Yang, Antoine Ameloot, Guillaume Lio, Angela Sirigu, Jean-René Duhamel

## Abstract

We used digit-tracking, a touch-based method for assessing visual attention, to investigate spontaneous exploration in macaque monkeys. By engaging with degraded images on a touch-sensitive display, monkeys could uncover high-resolution portions through finger movements, allowing for natural and unrestricted interaction. Monkeys received juice rewards after touching a predetermined number of pixels, but no specific regions were targeted. Attention maps were generated from their interactions, along with data from human digit-tracking and monkey eye-tracking experiments. We applied a saliency model and a Convolutional Neural Network (CNN) model to predict the empirical explorations. The correlation between model prediction maps and empirical attention maps indicated that monkeys focused non-randomly on information-rich regions, with the CNN model providing the most accurate predictions. These findings suggest that exploration was driven by intrinsic curiosity, beyond the extrinsic rewards for interaction. Digit-tracking offers a minimally invasive, portable alternative to eye-tracking, expanding research opportunities in visual cognition within ecologically valid settings.

## Introduction

Studies in non-human primates have significantly advanced our understanding of the mechanisms underlying visual attention in humans, largely due to the remarkable similarities in the organization of visual perception and cognition across species. These similarities enable researchers to draw insights from primate behavior to inform our understanding of human visual processing, including how both monkeys and humans explore visual images and prioritize relevant information. In both species, exploration tends to focus on visually salient regions^1,2^, such as the relevant features of objects^3^, body parts, and faces, which are key to survival and social interaction^4–6^. This preference for faces can be modulated by factors such as sex, rank and social perception features of the face^7–10^. Understanding how primates attend to and process visual stimuli thus offers valuable perspectives on the neural and cognitive mechanisms that govern attention in humans.

Traditionally, studies investigating visual attention in non-human primates have relied on eye-tracking methods. Although eye-tracking provides precise measures of visual exploration, these methods are often constrained by the need for extensive training^11–13^ and head restraint which can limit the ecological validity of the data. Additionally, the invasive nature of the equipment and the required calibration procedures can restrict the flexibility and portability of such studies.

Digit-tracking, a method wherein subjects explore images by using their fingers on a touch-sensitive display, presents a promising alternative to traditional eye-tracking^14^. By allowing for spontaneous and natural exploration without the need for head restraint or eye movement calibration, digit-tracking eliminates many of the constraints associated with eye-tracking studies. In this method, a Gaussian-blurred image is presented on a touch-sensitive interface and can be locally unblurred by sliding a finger over it. The blur simulates the low resolution of the peripheral retina, while the clear area revealed by touching the screen models the high foveal acuity, and eye movements are inferred from finger movements. This method has already been validated in human subjects, where it has proven to be an effective and minimally invasive technique for studying visual attention^14–16^. However, its application in non-human primates has yet to be thoroughly explored.

In this study, we aimed to test the feasibility and validity of digit-tracking in a group of laboratory macaques. We hypothesized that monkeys engage in visual exploration driven by intrinsic curiosity and predicted that they will demonstrate non-random, spontaneous exploration patterns by focusing on visually informative regions of the display, despite the absence of explicit cues or instructions to do so. We further hypothesized that digit-tracking exploration patterns would correlate with established metrics of visual attention and that a Convolutional Neural Network could predict these patterns based on the image features.

## Results

We trained five macaques to freely view 59 images – featuring animals, humans, objects and landscapes - using this digital interface within their home cage. For validation, we tested nine human subjects with the digit-tracking method and two of the macaques with standard eye-tracking, all using the same images (Figure 1). We began by evaluating the training procedure through an analysis of each monkey’s performance and generated attention maps to visualize their exploration patterns. We then used a standard saliency model and a Convolutional Neural Network model to predict the empirical attention maps including monkey digit-tracking, human digit-tracking, and monkey eye tracking.

**Figure 1.**
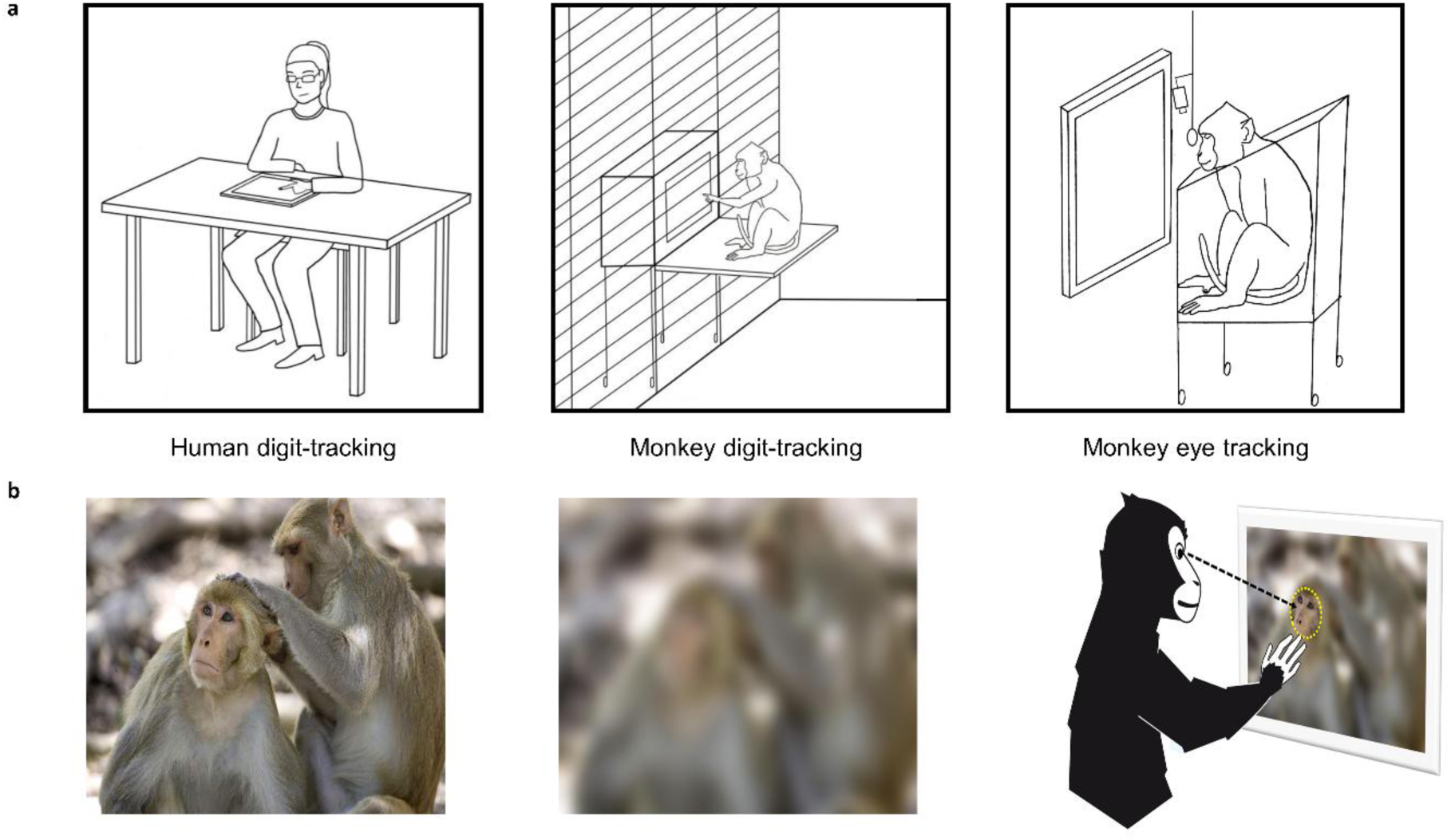
An illustration of the experimental conditions. **a)** demonstrates human digit-tracking, macaque digit-tracking, and macaque eye tracking experiments; **b)** shows an example of the original image, the Gaussian-blurred image and the blurred image with a clear aperture.

### Monkeys were engaged in the free-viewing task

Monkeys were trained to explore images on a touchscreen integrated into a mobile, battery-powered device equipped with a liquid delivery system. The device was positioned directly against their home cages, allowing them to perform the task in a low-constraint, familiar environment. Monkeys were not incentivized to touch specific areas. They could touch any location on the screen and a reward was automatically delivered after a predetermined number of pixels had been touched. During the training phase, the cumulative exploration distance required to complete a trial— measured as the distance traveled by the monkey’s finger across the screen—was progressively increased to a final criterion of approximately 56° of visual angle. On average, it took 9 training sessions for the monkeys to reach this final criterion and complete the entire image set within a single session, with individual variations ranging from 7 to 11 sessions. To determine if the monkeys were genuinely interested in the images, we computed the proportion of their exploration time on image. The images didn’t occupy the entire surface of the touchscreen, if their finger mainly landed outside the image boundaries, it indicated that they had developed a strategy to obtain the reward by touching the screen without actually paying attention to the image. Figure 2a showed that three of the monkeys mainly focused on the image contents, especially images of creatures such as monkeys, humans, dogs. To investigate whether their focus on the image content improved over sessions, we selected the first and last third of the sessions (floor the value if it wasn’t an integer) as early and late sessions, and performed a two-way ANOVA (Subject*Session). We observed an increase in the proportion of time spent on image over sessions, *F*(1,290) = 96.944, p < 0.001, 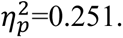 Moreover, post hoc results indicated that Monkey S (t(57) = 15.172, *p_tukey_* < 0.001, 95% CI [0.199, 0.309], d = 1.742) and Monkey J (t(57) = 3.634, *p*_*tukey*_ = 0.012, 95% CI [0.006, 0.116], Cohen’s d = 0.417) showed significant improvement over sessions (Figure 2b).

**Figure 2.**
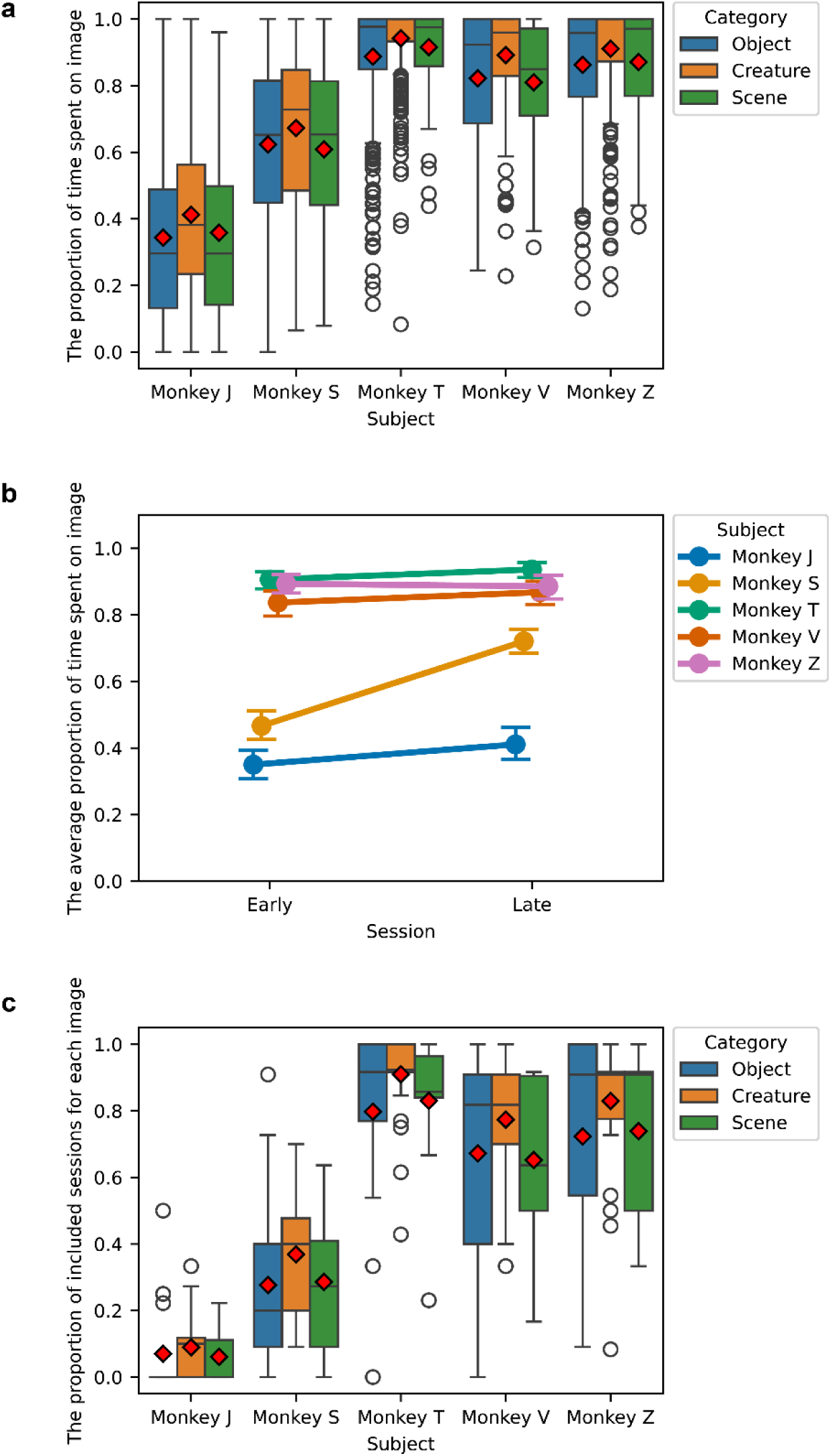
The inclusion of data based on the proportion of time spent on images. **a)** The boxplot of proportion of time spent on images of different subjects, the diamonds represent mean values of each condition; **b)** The change of mean proportion of time spent on images between early and late sessions; **c)** The boxplot of proportion of included sessions per image after filtering.

Subsequent analyses were performed on the attention maps derived from finger trajectories which showed the allocation of attention during exploration, so we deemed reasonable to exclude trials where the subjects did not exhibit sufficient task engagement, defined as touches within the image boundaries. We set the inclusion criteria for the proportion of time spent on the image at 0.8. Then, we computed the proportion of included sessions for each image. Figure 2c shows the proportion of included session for each image and subject. We found that only a few sessions from Monkey J and Monkey S met the inclusion criteria, which was insufficient for future analysis. Therefore, we excluded these two subjects. After excluding certain images and subjects, the average number of included sessions for the remaining subjects was as follows: Monkey T had an average of 11.138 sessions (SD=2.267), Monkey V had an average of 8.121 sessions (SD=2.760), and Monkey Z had an average of 8.678 sessions (SD=2.956). We also excluded one image because it only had one session from one single subject after exclusions.

### Characterization of attention maps

To present the exploration patterns, we generated empirical attention maps for each trial and computed the average attention maps across subjects and sessions. Figure 3a shows the average attention maps from macaque digit-tracking for five different images, and figure 3b shows the attention maps for one image from different data sources. We observed that the explorations were guided by objects, contrasts, features and social relevance. Notably, when viewing images containing living creatures, the monkeys tended to spend more time on faces, which is consistent with many previous eye-tracking studies^4–6^. The exploration patterns showed similarities across different data sources, which we will quantify in the next section.

**Figure 3.**
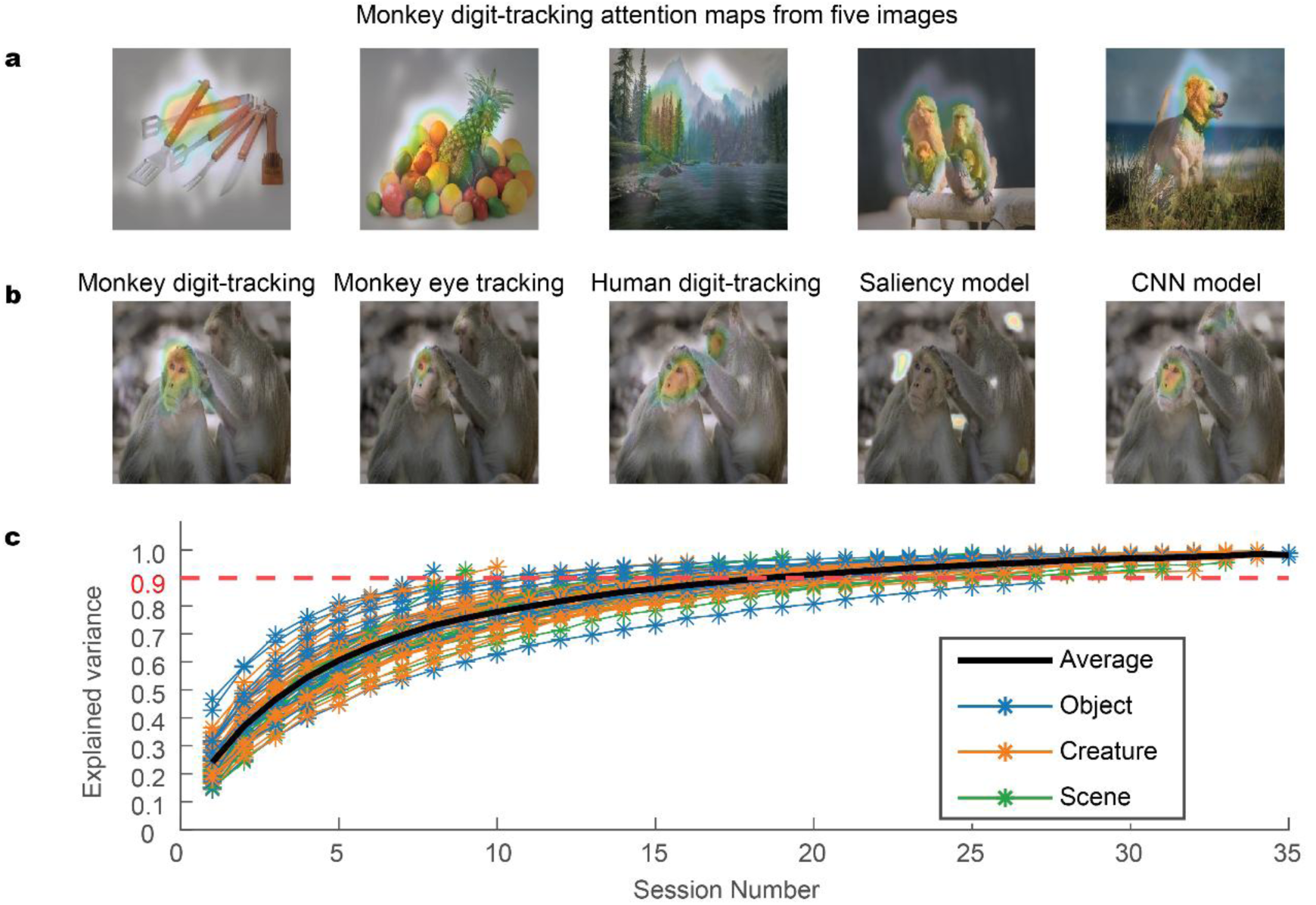
Characterizations of attention maps. **a)** shows attention maps of five different images from the monkey digit-tracking data and **b)** shows attention maps of the same image collected from different experiments and the prediction maps from two models; **c)** The curves of explained variance against number of sessions.

**Figure 4.**
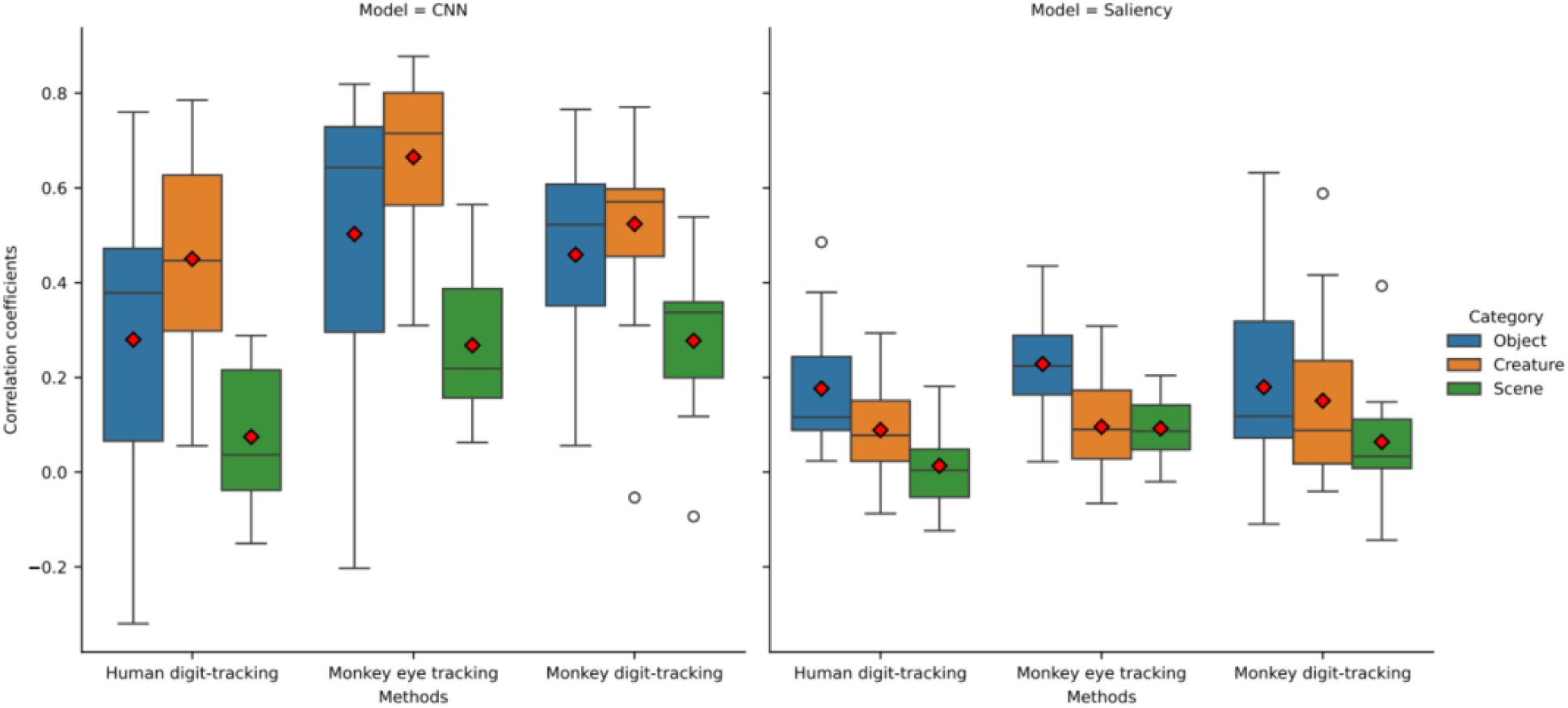
The boxplots of the correlation coefficients between the model prediction maps and the empirical attention maps (the red diamonds indicate the mean values). The CNN model predicts the exploration better than the saliency model. Its performance in creature images is the best while the performance of saliency model in the object images is the best.

To determine the number of sessions needed for reliable measurements, we performed a convergence analysis as follows: For each image, we (1) generated average attention maps using incrementally larger session samples (from N=1 to maximum available), (2) calculated the correlation between these sample-based maps and the ground-truth map (derived from all available sessions), and (3) plotted the explained variance (R²) against session count (Figure 3c).The analysis revealed that an average of 17.69 sessions (SD = 4.09) were required to achieve 90% of the maximum explainable variance. This estimate was robust across our dataset, with ≥18 sessions available for analysis from at least 3 subjects on 51 different images.

### Digit-tracking captures features predicted by models

To assess whether digit tracking reflects genuine attentional allocation, we compared empirical attention maps - derived from monkey digit-tracking, human digit-tracking, and monkey eye-tracking – against predictions from two computational models. Critically, rather than directly comparing the three empirical datasets (which differ in species and modality), we evaluated how well each aligned with model-generated attention maps. This indirect approach allowed us to test whether digit-tracking captures similar attentional biases as eye-tracking and whether these biases are driven by shared visual features across species. While this approach does not quantify species- or modality-specific differences directly, it provides a unified framework to evaluate attentional commonalities across datasets. We selected two models with distinct strengths: (1) a saliency model (Walther and Koch^17^), which computes low-level visual features to predict bottom-up attention, (2) a Convolutional Neural Network (CNN) ( Lio et al^14^), trained on human digit-tracking and eye tracking data, which incorporates both low-level saliency and higher-level social cues (see Methods for details of the two models). A two-way ANOVA (Model * Method) revealed that CNN’s prediction maps correlated significantly better with empirical attention maps than the saliency model (*F*(1,58) = 119.375, *p* < 0.001, 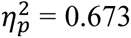), consistent with the presence of social content in our stimuli that the saliency model could not capture.

To evaluate model performance across methods and stimulus categories, we conducted separate two-way ANOVAs (Method*Category) for each model. For the CNN model, significant main effects were observed for both method (*F*(2,112) = 37.887, *p* < 0.001, 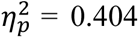) and category (*F*(2,112) = 12.530, *p* < 0.001, 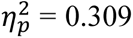), with a non-significant interaction (*F*(4,112) = 2.178, *p* = 0.076, 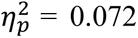). Post hoc comparisons revealed that prediction accuracy was highest for human digit-tracking significantly surpassing both monkey digit-tracking (t(56) = 8.721, *p*_*bonf*_ < 0.001, 95% CI [0.151, 0.270], d = 0.967) and monkey eye tracking (t(56) = 5.563, *p*_*bonf*_ < 0.001, 95% CI [0.027, 0.085], d = 0.699). Monkey eye-tracking predictions were also more accurate than monkey digit-tracking, t(56) = 2.512, *p*_*bonf*_ = 0.045, 95% CI [0.001, 0.116], d = 0.268. Regarding stimulus categories, the CNN showed stronger correlations for creature (t(56) = 4.963, *p*_*bonf*_ < 0.001, 95% CI [0.171, 0.509], d = 1.563) and object images (t(56) = 2.908, *p*_*bonf*_ = 0.014, 95% CI [0.031, 0.383], d = 0.953) compared to scenes.

The saliency model yielded different results, with weaker but still significant main effects for method (*F*(2,112) = 3.649, *p* = 0.029, 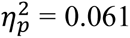) and category (*F*(2,112) = 6.930, *p* = 0.002, 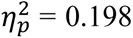), and no significant interaction (*F*(4,112) = 2.282, *p* = 0.065, 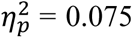). Unlike the CNN, the saliency model showed no significant difference between monkey digit-tracking and eye-tracking (t(56) = 1.901, *p*_*bonf*_= 0.187, 95% CI [0.011, 0.088], d = 0.295), though human digit-tracking predictions remained more accurate than monkey digit-tracking (t(56) = 2.863, *p*_*bonf*_= 0.018, 95% CI [0.006, 0.085], d = 0.352). For stimulus category, the saliency model performed best with object images, outperforming both creatures (t(56) = 2.687, *p*_*bonf*_ = 0.028, 95% CI [0.007, 0.159], d = 0.638) and scenes (t(56) = 3.496, *p*_*bonf*_ = 0.003, 95% CI [0.041, 0.236], d = 1.062).

To complement our correlation-based findings, we examined the spatial precision of the models by measuring distances between the most salient point in model predictions and empirical attention maps. This alternative metric revealed a marginally significant main advantage for the CNN model overall (*F*(1,58) = 4.381, *p* = 0.041, 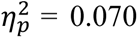). Subsequent separate ANOVAs (Method × Category) showed, for the CNN model, a marginally significant main effect of category (*F*(2,112) = 3.223, *p* = 0.047, 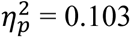) and a significant interaction effect (*F*(4,112) = 3.219, *p* = 0.015, 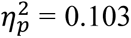), though post hoc tests did not reach significance. As shown in Figure 5, the CNN’s predicted salient points tended to be closer to empirical data for creature images than other categories. For the saliency model, no significant effects emerged (method: *F*(2,112) = 0.791, *p* = 0.456, 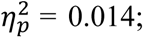 category: (*F*(2,112) = 0.318, *p* = 0.729, 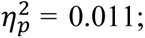 interaction: *F*(4,112) = 1.759, *p* = 0.142, 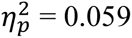).

**Figure 5.**
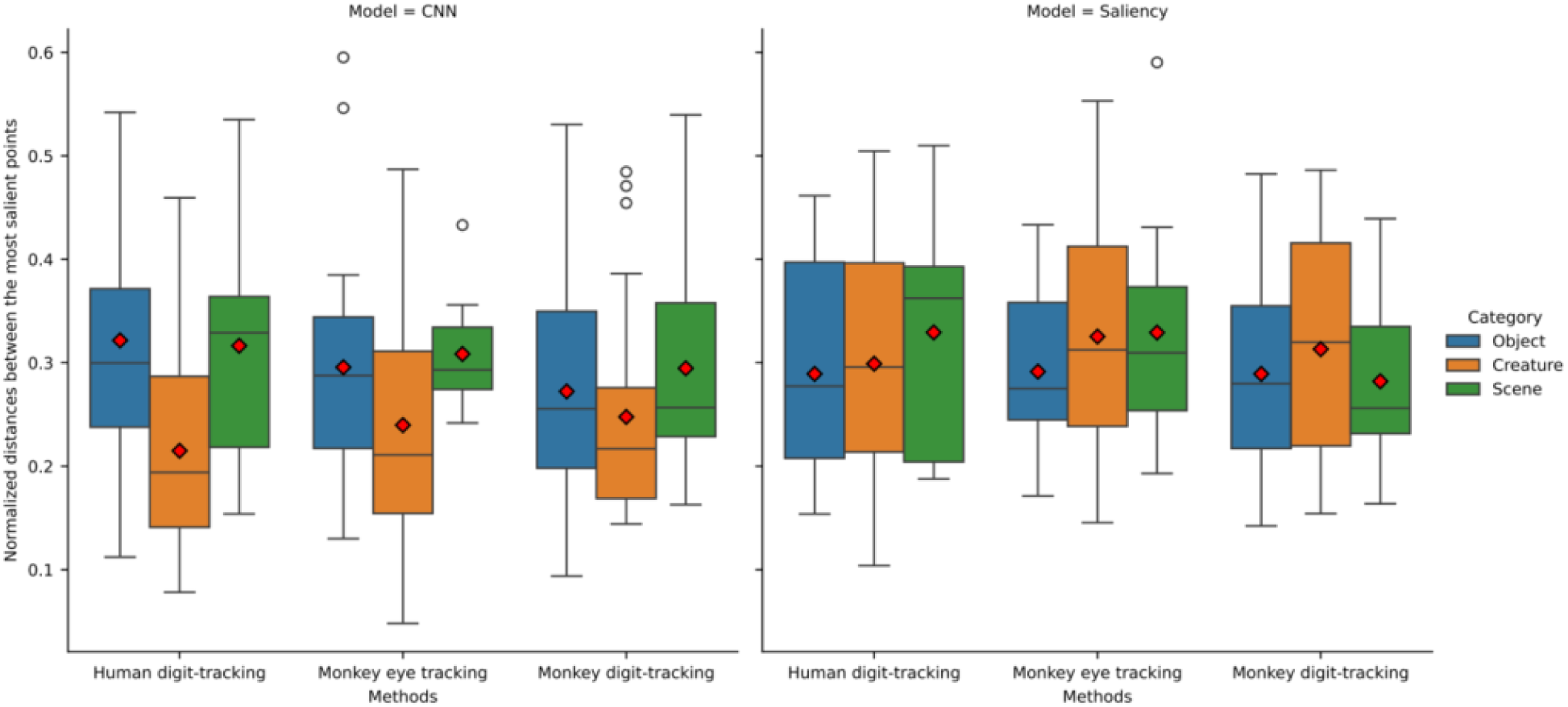
The boxplots of the normalized distance between the most salient points (the red diamonds indicate the mean values). For CNN model, the distance between prediction maps and the attention maps in scene images are slightly shorter even the test was not significant.

When combining results from both correlation and distance-based metrics, the CNN model demonstrated superior performance, particularly for human data and socially relevant stimuli. This advantage may reflect its training on human-derived data and capacity to integrate social cues. In contrast, the saliency model showed limited predictive power, with its only robust performance occurring only for object images in correlation analyses. Together, these results quantitatively establish that the CNN better captures attentional patterns across exploration methods, with its performance advantage most pronounced for socially relevant content.

### Shared feature selection in human and macaque visual exploration

Building on our previous CNN architecture developed in Lio et al.^14^, we investigated whether humans and macaques attend to similar visual features during exploration. While the previous section’s analyses used the pre-trained model from that study, we extended this approach by retraining the model on our current dataset to compare feature weighting across species and modalities. In our original implementation^14^, we adapted a well-established Convolutional Neural Network (CNN) architecture (AlexNet Network^18^) to detect essential features during visual exploration. This architecture was initially trained for image classification using the ImageNet database (http://www.image-net.org) for the ImageNet Large-Scale Visual Recognition Challenge (ILSVRC^19^). We took the first five convolutional layers and created prediction maps by linearly combining their feature maps. The weights for this combination of feature maps were learned from human empirical data. For the current analysis, we retrained the model on our new dataset to directly compare how features are weighted across different species (human vs macaques) and testing methods (digit-tracking vs eye-tracking). A key aspect of our analysis involved examining how image resolution affects feature detection, through what we refer to as the “pyramid level,” where higher levels correspond to finer image analysis. By manipulating this level, we can examine how the model’s ability to detect and utilize features changes with varying degrees of image detail.

The results revealed three important findings. First, the relative importance (weighting) of the 256 features in our new analysis closely matched the patterns we previously reported^14^ (Figure 6a). Second, when we compared how features were weighted between humans and monkeys using digit-tracking, we found remarkably strong correlations at all pyramid levels (0.96, 0.94, and 0.94 for levels 1-3 respectively). Third, the correspondence between monkey eye tracking and digit-tracking weights became slightly weaker at finer resolutions (0.84 at level 1 to 0.77 at level 3). This pattern suggests that while humans and macaques fundamentally attend to similar visual features during exploration, digit-tracking may capture these features at a slightly coarser resolution compared to eye-tracking particularly when examining fine details. The high correlations across species indicate shared mechanisms of feature selection, while the resolution-dependent differences highlight how measurement method can affect the precision of captured attentional patterns.

**Figure 6.**
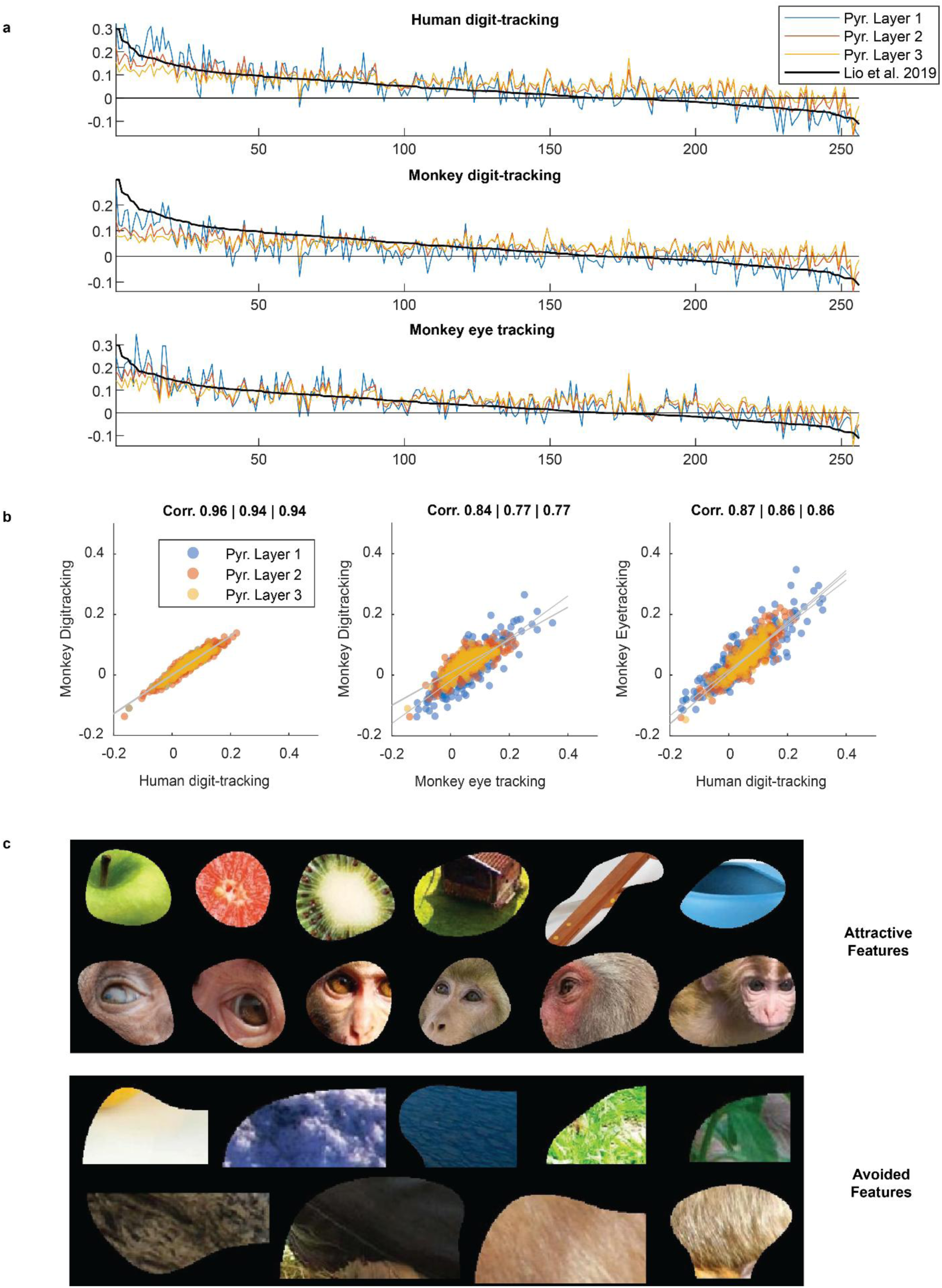
The features decomposed by the Convolutional Neural Network architecture. a) The hierarchical orderings of the features from each testing condition. b) shows the scatter plot of feature orderings. c) shows examples of attractive and avoided features shared by human and macaques. Both species were attracted by salient fruits, objects, eyes and faces, whereas they were not interested in the background and peripheral areas.

## Discussion

Digit-tracking, initially designed for human subjects, has proven to be a reliable and innovative tool for studying spontaneous attention allocation in non-human primates in this study. One significant advantage of digit-tracking is its non-invasive nature, which allows data collection in a more relaxed, free-ranging environment, minimizing the stress often associated with traditional laboratory-based methods. We successfully trained five macaques to explore the Gaussian-blurred images on a touch screen within their own cages, achieving stable performance from three of them. The empirical attention maps suggested that the explorations were guided by image characteristics and could be predicted by a Convolutional Neural Network (CNN) model. The analysis of monkey eye tracking and human digit-tracking data revealed similar pattern as monkey digit-tracking, with the most accurate predictions occurring for creature images. The decomposition of the exploration pattern using the CNN model further demonstrated that human and macaques are attracted to and avoid similar features. On average, 18 sessions were required to reach stable and reproducible performance, which is not particularly difficult to achieve under this experimental setting. This suggests that digit-tracking can provide a reliable method for studying attention allocation in non-human primates.

In this study, we collected digit-tracking data while the subjects were in their cage. They were not forced to always stay in front of the screen but had to reach a predetermined exploration distance to complete each trial. During training, we gradually increased the exploration distance to reach the predetermined value. The subjects were relaxed and showed no signs of anxiety or resistance to the equipment. All subjects adapted to the task quickly, including some monkeys who had a history of being difficult to train in a laboratory setting. We collected good-quality data from three of the five subjects. Monkey S improved his concentration on the images over sessions and produced satisfactory results. In future studies, an improvement could involve estimating the proportion of time spent viewing the image during the training stage to identify uncooperative subjects. This would allow us to exclude subjects like Monkey J, who did not engage with the task, before the formal experiment. In another study (unpublished), we tested an alternative method for ending the trial. The images were presented for a fixed duration triggered by the first contact and then the subject got the reward regardless of exploration distance accomplished. We didn’t use this method in current study because we were uncertain whether it would yield a sufficient amount of exploration data. However, we observed that the subjects continued to interact with the screen even under the fixed duration condition, suggesting that these actions were driven by genuine intrinsic motivation to acquire visual information. This method may offer an alternative approach for future experiments.

Our attention allocation analyses revealed striking cross-species and cross-method similarities. The CNN model’s consistent superiority over the saliency model – particularly for social stimuli (faces, eyes, and bodies of conspecifics) confirm that macaque attention, like human attention, prioritizes meaningful features beyond raw visual salience^20–22^. Although the CNN showed slightly reduced accuracy for monkey digit-tracking versus eye-tracking, the preserved hierarchical orderings of features across all conditions suggests this primarily reflects measurement precision differences rather than distinct attentional strategies. Most importantly, the conserved feature importance patterns imply an evolutionarily shared attention mechanism that operates similarly whether exploration occurs via fingers or eyes.

In conclusion, our findings demonstrate that digit-tracking can be a valuable, adaptable tool for studying spontaneous visual behavior in non-human primates. Monkeys explored information-rich regions of the image display, in a manner consistent with established metrics of visual attention. This research contributes to a broader understanding of attention mechanisms in primates, showing that visual exploration involves an intrinsic component related to curiosity. Our findings open up new avenues for research on the cognitive and motivational foundations of attention, as well as practical applications for studying primate behavior outside the constraints of traditional lab environments. By demonstrating a method that could ultimately benefit a range of animal research, from controlled lab studies to free-ranging wildlife observation, this study introduces digit-tracking as a flexible, scalable, and insightful approach for primate visual cognition research.

## Methods

### Participants

For this study, we collected data from five macaques at ISC-MJ, including two rhesus macaques (Monkey V, male, 11.2 kg, 11 years old; Monkey Z, female, 8.8 kg, 19 years old) and three long-tailed macaques (Monkey T, male, 9.5 kg, 12 years old; Monkey S, 9.7 kg, 12 years old; Monkey J, 9.9 kg, 12 years old). In addition, we obtained the eye tracking data from Monkey V and Monkey T.

Animals were provided with a diet consisting of monkey chow, fresh fruits, and vegetables, and their water intake was regulated. Each morning, the monkeys received a minimum of 50 mL of water with their daily portion of monkey chow, and the task began shortly after the water and food were provided. During the task, they received approximately 0.5 mL of water for each successfully completed trial. The total daily water allowance was adjusted based on each monkey’s performance: if a monkey did not complete the entire set of images in a single session, the water provided after the task was gradually reduced until they consistently completed the full set in one session, after which the acquisition phase was initiated. Throughout the study, the minimum daily water intake for each animal was maintained at no less than 20 mL per kilogram of body weight, regardless of task performance, to ensure proper hydration and overall health. Finally, each monkey had one day of free access to water each week. Enrichments were offered several times a week with different toys or substrates (boxes and puzzles containing dry fruit, swings) that promote play, curiosity, object manipulation, and foraging following recommendations from our own laboratory animal welfare committee.

A group of nine human participants (mean age = 27, SD = 5.7) with no history of psychiatric or neurological disease were recruited from the pool of graduate students and technician of the Institute of Cognitive Science Marc Jeannerod of Bron (ISC-MJ), France (FR).

### Picture database

A total of 59 images depicting various aspects of natural and social scenes, including humans, monkeys, other animals, landscapes, objects, and abstract art, were selected. The selection of images for the database intentionally encompassed a wide range of content to ensure that it did not limit the exploratory behavior and maintained the curiosity of both the tested human and monkey subjects. The picture database includes images featuring the bodies and faces of both humans and monkeys because these stimuli elicit distinctive exploration patterns that serve as an ideal basis for conducting comparative analyses across different species and contexts. Images were categorized into three groups: object (20), scene (12) and creature (27).

### Digit-tracking method

The digit-tracking method was implemented following the same framework described in Lio et al^14^. Stimulus presentation, real-time picture modification and data recording were written using the Psychophysics Toolbox for MATLAB (R2015a, the MathWorks, Inc.).

This method consists into presenting on a capacitive touchscreen display a blurred image simulating an approximation of the acuity present in peripheral vision. When the subject touched the screen with its finger, an area approximating the size of the fovea (see details below) was displayed at full resolution just above the point of contact. The area was shifted upwards to prevent the subject’s view masking by the finger. Removing the finger from the display restored the image’s fully blurred appearance and sliding the finger across the display caused the full resolution area to be updated. Coordinates of the area’s center, regarded as the proxy for eye positions and attention, were continuously recorded.

In the implementation of the digit tracking method, three critical parameters need to be accounted for: the degree of picture degradation to simulate an approximation of peripheral vision, the dimensions of the foveal area being simulated, and the distance of the foveal aperture relative to the point of contact. We applied a Gaussian blur filter with a size parameter of σ=1.35° to simulate a more pronounced blur compared to standard peripheral vision as it has previously demonstrated that slightly higher blur level promotes image exploration^15^. Additionally, given that the monkeys tested in their home cage with the digit-tracking method were not head-fixed, the distance between their eyes and the screen could not be controlled precisely. However, this distance could not be less than 16.5cm, as it corresponded to the distance between the screen and the bars of the home cage. Therefore, applying a more intense blur guaranteed that the image resolution remained below the acuity of peripheral vision, even under the most minimal eyes-screen distance of 16.5cm. A Gaussian aperture window with a size parameter of σ=2.6° was used to simulated foveal vision. The aperture window size was slightly enlarged compared to that used in the original study^14^ to encourage exploration in monkeys. The distance between the point of contact and the center of the foveal vision was 1.4°.

### Eye tracking method

As subjects V and T were taking part in electrophysiology experiments of other studies, these individuals were equipped with head-restraint device that facilitated the recording of their eye positions using an infrared eye tracker (ISCAN ETL200 Series, sampling rate = 240 Hz) while they explored the images. The open source software NIMH MonkeyLogic was used for task control and data acquisition^23^. Each session begins with a calibration routine of 9 points using a fixation task. Throughout the experiment, the calibration’s stability was continuously monitored by the operator, and if any deviation or inconsistency was detected, the calibration process was repeated to ensure accurate and reliable eye-tracking data.

### Apparatus and procedures

The experiment consisted in recording the exploration of pictures by both monkeys and humans. Monkeys’ exploration was tracked using either the digit-tracking method in their home cage or eye-tracking in laboratory. For humans, only the digit-tracking method was used.

The task structure remained consistent across all conditions, but subjects interacted with the screen differently depending on whether they were tested using the digit-tracking or eye-tracking methods. In the digit-tracking condition, each trial began with a green virtual button appearing at the center of the screen against a black background. The subject initiated the presentation of the blurred image by touching this button. The subject could then freely explore the image until a track-path of a predetermined length has been reached. The image was then removed and monkey subjects were rewarded with a drop of juice. After a fixed 2-second delay to allow the reward to be consumed, the next trial began. Trials continued until all images of the database had been explored, with the order of their presentation following a pseudo-random sequence. All monkeys underwent identical familiarization and training procedures as follows: in the beginning, subjects were only required to touch the green virtual button to receive a reward until they became accustomed to interacting with the touchscreen, typically within fewer than 2 sessions. Subsequently, blurred images were introduced upon button touch, and the track-path to reach trial completion was gradually extended, starting from the minimum cumulative exploration distance (requiring only one contact with the image) and progressing to the final distance to reach. The daily cumulative distance increment was adjusted for each monkey based on their performance. On average, it took 9 sessions to complete the 59 trials with the final cumulative distance (9 sessions for monkey V, 10 sessions for monkey T, 11 sessions for monkey Z, 8 sessions for Monkey J and 7 sessions for Monkey S).

To test subjects in their home cage with the digit-tracking method, a tablet computer (Dell Latitude 7210, frequency of 60 Hz, 12.3″ in diagonal, resolution of 1920 x 1280 pixels) was integrated into a mobile, battery powered training box equipped with a liquid delivery system and a seating platform. This mobile device was positioned against the bars of the subject’s cage, providing access to the touchscreen to perform the task. The cumulative exploration distance required to complete a trial was adapted for monkeys and set at 12 inches (2250 pixels with a screen resolution of Pixels Per inch = 188, corresponding to ≈ 56° of visual angle) to prevent disengagement and ensure their ability to complete the trials.

In the eye-tracking condition, monkeys were seated in a primate chair facing a capacitive touchscreen (ELO® 2770L, frequency of 60 Hz, 27″ in diagonal, resolution 1920 × 1280 pixels). Their heads were immobilized to enable eye-tracking, as described previously. The dimensions of the images were adjusted to ensure that their sizes in visual angles were similar to those used in the macaque digit-tracking condition. The structure of the task was similar to that of the digit-tracking condition, except that image presentation was initiated by fixation of a green target for 500 milliseconds. To ensure comparable exploration in both the macaque digit-tracking and eye-tracking experiments, the images were displayed for 2 seconds. This duration corresponds to the average time necessary to achieve 56° of visual angle exploration, which is the cumulative distance needed to complete a trial in the digit-tracking experiment. Subjects were required to keep their eye position within the screen limits during the trials. Exceeding these limits terminated the trial without reward delivery, and the image was presented again later in the session. Subjects underwent training until they achieved a performance level of 80% successful trials before beginning formal data acquisition.

In the human digit-tracking condition, participants were instructed to sit in front of the same tablet computer used in the macaque digit-tracking condition (Dell Latitude 7210, frequency of 60 Hz, 12.3″ in diagonal, resolution of 1920 x 1280 pixels) without any constraint. Each participant was instructed to freely explore each picture during a single session lasting approximately 30 min. The picture was removed when the cumulative distance of exploration reached 22.3 inches (4000 pixels with a screen resolution of Pixels Per inch = 188). The selection of this distance was guided by participant feedback collected in a preliminary pilot study to enable sufficient image exploration without inducing disinterest to maintain participants’ engagement throughout the task.

### Data analyses

All the analyses were conducted in MATLAB (R2015a, the MathWorks, Inc.) using our custom scripts. Some plots were generated in Python (3.9.19) using the Seaborn library (0.13.12). Statistical tests were conducted in JASP^24,25^.

### Attention maps

Attention heat maps were computed as described in Lio et al^14^. Briefly, coordinates of tracking points were transformed from the screen space to the picture space. Attention map for each subject and each picture, i.e. the probability density of exploration, was calculated from the tracking data using kernel density estimation with a Gaussian kernel weighted by duration and a bandwidth of 1° of visual angle. Each subject-level density was then normalized using the Min-Max normalization. Finally, group-level maps were generated by averaging scaled subject-level densities and normalized using the same normalization procedure. Attention maps for eye-tracking data were computed following the same procedure based on fixations.

### Convergence analysis

For each image and each iteration, we generate a random permutation of session numbers of this image. Then we generated attention maps from 1 to N-1 sessions (N = number of sessions for each image) and computed the correlation between these attention maps and the average attention map across all sessions. This procedure was repeated 40 times and then we plotted the curves of the mean coefficient of determination over sessions. We set the criteria of explanation variance as 0.9 and compute the number of sessions needed to reach this criterion.

### The saliency maps

The saliency maps were generated using the approach described by Walther and Koch^17^, which is a variation of the classical saliency model originally proposed by Itti and Koch^26^. The authors developed a Saliency Toolbox written in MATLAB and made it available on Github (https://github.com/DirkBWalther/SaliencyToolbox). We utilized this toolbox with its default parameters to produce the saliency maps.

### The Convolutional Neural Network model

The Convolutional Neural Network model was trained by Lio et al^14^. We utilized the pre-trained AlexNet Network^18^ to generate prediction maps. The selected model was trained on a subset of more than one million images, extracted from the ImageNet database (http://www.image-net.org) for the ImageNet Large-Scale Visual Recognition Challenge (ILSVRC^19^). Dedicated originally for large-scale image classification, the CNN architecture was transformed using the Matlab Neural-Network toolbox (Mathwork Inc.) to solve a regression problem and generate predictions. First, the outputs of the fifth convolutional layer, rectified by a linear unit (ReLU layer), were interpolated to obtain 256 feature maps with the same resolution of the input image (227 × 227 pixels). Then the maps were linearly combined using wCorr, a fast estimate of the weights by averaging the Pearson Correlation Coefficients calculated between a feature-map and the measured saliency map (the 256 weights are learned independently). This approach can be considered as a simplified implementation of the Kümmerer et al ^27^. method for saliency map prediction.

### The distances between most salient points

First, we identified the most salient point (i.e., the pixel with the highest value) in each map. For each image, we had one prediction map from the saliency model and one prediction map from the CNN model, but multiple empirical attention maps across different subjects or sessions. We then measured the distance between the most salient point in each model’s prediction map and the most salient point in each empirical attention map, normalized by dividing by the diagonal length of the image. Finally, we averaged these distances for each image within each recording method.

## Code availability

The example code and data are available on the Open Science Framework (https://osf.io/na4mv/)

## Ethics approval

Animal experiments conducted at the ISC-MJ (User Establishment n°69 029 0401) were authorized by the local Animal Ethics Committee (permit n°2021031218457217). Human experiments were approved by Ethics Committee O5 – Ouest V (IDRCB 2019-A02160-57, project N° 19.07.30.45727) and a written informed consent was obtained from all participants.

## Conflicts of interests

All authors declare no conflict of interests.

## Author information

These authors contributed equally: Y.Y and A.A. Y.Y, A.A and J.R.D designed the study; A.A collected the data; Y.Y analyzed the data and prepared the figures; G.L trained the Convolutional Neural Network models; Y.Y, A.A and J.R.D interpreted the data and wrote the manuscript; J.R.D supervised the project; All authors contributed to the writing of the manuscript.

## Acknowledgments

This work was supported by grant from the European Research Council (ERC) under the European Union’s Horizon 2020 research and innovation program (grant agreement no. 885746) to J.R.D. We wish to thank Marine Cuvilliez for the drawings of procedure illustrations and Fidji Francioli and the animal facility staff for the animal care.

## Notes

### Competing Interest Statement

The authors have declared no competing interest.

